# Timecourse and source localization of abstract and concrete semantic representations

**DOI:** 10.1101/2023.06.23.546231

**Authors:** Lorenzo Vignali, Yangwen Xu, Jacopo Turini, Olivier Collignon, Davide Crepaldi, Roberto Bottini

## Abstract

Dual coding theories of knowledge suggest that meaning is represented in the brain by a double code, which comprises language-derived representations in the Anterior Temporal Lobe and sensory-derived representations in perceptual and motor regions. This approach predicts that concrete semantic features should activate both codes, whereas abstract features rely exclusively on the linguistic code. Using magnetoencephalography (MEG), we adopted a temporally resolved multiple regression approach to identify the contribution of abstract and concrete semantic predictors to the underlying brain signal. Results evidenced early involvement of anterior-temporal and inferior-frontal brain areas in both abstract and concrete semantic information encoding. At later stages, occipito-temporal regions showed greater responses to concrete compared to abstract features. The present findings shed new light on the temporal dynamics of abstract and concrete semantic representations in the brain and suggest that the concreteness of words processed first with a transmodal/linguistic code, housed in frontotemporal brain systems, and only after with an imagistic/sensorimotor code in perceptual and motor regions.

## 1. Introduction

Abstract and concrete semantic representations form fundamental aspects of word meaning. For instance, concrete features (e.g., four legs, red fur), as well as abstract features (e.g., intelligent, aggressive), contribute to the representation of a fox in semantic knowledge (Borghesani & Piazza, 2017; Smith et al., 1974). Although the ability of the brain to retrieve these representations is at the core of human semantic knowledge, the neural underpinnings of this process are not completely understood.

To explain how various sorts of knowledge guide a broad range of behaviors, Dual Coding Theories (DCT; Paivio, 1986, 1991) put forward the idea that, in the brain, semantic knowledge is represented in a dual coding system comprising a linguistic code, and an imagistic/sensorimotor code. In DCT, abstract and concrete concepts can both be represented through the linguistic code, whereas the imagistic/sensorimotor code is available only for concrete aspects of meaning. In experimental psychology, the fact that concrete words are recognized faster (Kroll & Merves, 1986; Schwanenflugel et al., 1988; Schwanenflugel & Stowe, 1989) and memorized better (Allen & Hulme, 2006; de Groot, 1989; Fliessbach et al., 2006) than abstract words (i.e., the concreteness effect) has been traditionally taken as evidence in favor of the dual coding approach: only concrete concepts can activate both codes (linguistic and sensorimotor), a condition which gives them a processing advantage (Connell & Lynott, 2012; Paivio, 1986). However, the study of the concreteness effect in congenitally blind people has cast doubts on this interpretation (Bottini et al., 2021). Despite lacking a sensory code for visual features, early blind people processed visual unimodal-concrete words (e.g., “red,” “multicolor,” “transparent”) faster than abstract words, showing a concreteness effect that was indistinguishable from the one of sighted people (Bottini et al., 2021) and suggesting that the concreteness advantage is not driven by the availability of a double code for concrete words.

If, on the one hand, the study of blind people has shaken the confidence in psychological evidence considered a hallmark of dual coding models, on the other hand, it has revived the interest in DCTs from a neuro-cognitive perspective. Are sensory-derived and non-sensory-derived representations encoded in dissociable brain codes? A recent line of studies exploring the brain basis of visual knowledge in the absence of vision has provided alternative neurocognitive evidence for a dual code of semantic knowledge in the brain (see Bi, 2021). Two studies focusing on color representations in sighted and congenitally blind have shown that posterior brain areas in the V4 complex encode the similarity of color words, but only in sighted people. However, color similarity is also encoded in the dorsal anterior temporal lobe (ATL) in both sighted and blind (Bottini et al., 2020; X. Wang et al., 2020). Thus, the dorsal ATL seems to provide a non-sensory code to represent knowledge, both concrete and abstract, whereas a perceptual code for concrete representations relies on posterior perceptual regions and may not be available in the case of sensory deprivation (Bi, 2021).

Beyond research with populations devoid of specific aspects of perceptual experience, functional resonance imaging (fMRI) studies investigating topological as well as functional properties of the semantic network provide additional evidence in favor of a dual-code account of semantic knowledge in the brain (e.g., Bi, 2021; Xu et al., 2017). From a network perspective, the dorsal anterior temporal lobe (dATL) and posterior sensory and motor regions are components of dissociable brain systems. The dATL belongs to the high-level linguistic system in the left perisylvian network, encompassing the inferior frontal gyrus, the lateral temporal cortex, and the inferior parietal cortex (for instance, Fedorenko et al., 2011; Friederici, 2011). It has stronger connections to the other regions in the language network than the sensorimotor regions(Jackson et al., 2016; X. Wang et al., 2020). The left perisylvian language network is consistently activated in semantic tasks (Binder et al., 2009; Xu et al., 2017), and is considered to play a role in language-supported semantic processing (Xu et al., 2017). Beyond DCTs, other models suggest that both codes are present in the ATL, arranged in a continuous transmodal gradient (e.g., Lambon-Ralph et al., 2017). That is, the ATL is considered a transmodal/graded hub with a linguistic neural code in its dorsal part, and a perceptual code in its ventral part (Hoffman et al., 2015; Visser & Lambon Ralph, 2011).

On the contrary, visual regions (including the color region V4) belong to the highly distributed sensorimotor brain system, (see Wang et al., 2020) which reflects relevant perceptual dimensions of the input such as visual, tactile, auditory, etc. (Barsalou et al., 2003; Binder et al., 2005, 2009; Binder & Desai, 2011; Hoffman et al., 2015; Kana et al., 2012; Sabsevitz et al., 2005). These regions are usually more active for concrete compared to abstract concepts (Binder et al., 2005, 2009; Binder & Desai, 2011; J. Wang et al., 2010) and may host sensorimotor simulations of perceptual referents during semantic processing. However, as fMRI suffers from poor temporal resolution, several questions about the spatiotemporal dynamics of the dual code of knowledge in the brain remain unanswered. For instance, it is unclear whether transmodal/language-derived representations in the ATL are activated before, after, or at the same time as sensorimotor representations in perceptual regions. This missing information is crucial to understand the neural dynamics of conceptual processing and, in particular, how concreteness (abstractness) is encoded in the brain.

To answer these questions, we took advantage of the high temporal resolution of magnetoencephalography (MEG) signals combined with source-reconstruction techniques to assess the spatiotemporal dynamics of abstract and concrete semantic representations. Forty-six participants performed a semantic categorization task on 438 written words. Each word referred to a concept (e.g., chair, dog, policeman) that was independently rated across 65 feature dimensions (e.g., color, shape, happiness, arousal, cognition, etc.; Binder et al. 2016). Using principal component analysis (PCA), we reduced the dimensionality of this feature space into one abstract and one concrete semantic principal component. We then used a combination of multiple linear regression analysis and source reconstruction methods to assess neural dynamics of abstract and concrete semantic representations while keeping into account other types of psycholinguistic information processed during visual word recognition.

## 2. Material and Methods

### 2.1. Participants

Forty-six native Italian speakers (29 female, aged 24.8 ± 4.2 years) participated in the study. All participants were right-handed and had no history of neurological or psychiatric disorders. Before testing, participants gave their written informed consent and received monetary reimbursement for their participation. The experiments were conducted in accordance with the Declaration of Helsinki and were approved by the local ethical committee of the University of Trento.

### 2.2. Experimental design

We derived our stimulus set from a previous work by Binder and colleagues (Binder et al., 2016). Out of 535 English words filed in Binder et al.’s (2016) original work, 438 were translated into Italian (352 nouns in the singular form, 54 verbs in the infinite tense, and 32 adjectives in the singular masculine form). Selected words could be unambiguously translated into Italian. Participants were instructed to categorize each stimulus as either related to sensory-perception (i.e., they express something that is related to one or more of the senses), or unrelated to sensory perception. Visual stimuli were projected on a translucent whiteboard (1440×1080 pixel resolution) using a ProPixx DLP projector (VPixxTechnologies, Canada) at a 120 Hz refresh rate. Stimulus presentation was controlled via Psychtoolbox (Kleiner et al., 2007) running in a MATLAB 2015a environment. At the beginning of each trial, a 1s blank screen followed by a 0.5s fixation cross preceded stimulus appearance. Words appeared in a white monospaced bold font on a dark gray background, covering on average 3.2 degrees of visual angle (*SD* = 0.8). Stimuli remained on the screen for 0.3s, followed by a 1.7s blank screen. After this delay, a text (“Was it a word related to the senses? YES - NO”) prompted participants’ responses via button press operated with the dominant hand’s index and middle fingers. The response mapping was counterbalanced across participants. The maximum time given to respond was set to 2s and was followed by an interstimulus interval randomly jittered between 0.3s and 0.6s. Participants were familiarized with a short version of the task (30 trials taken from a different stimulus set) on a portable PC outside the MEG chamber. Each testing session lasted approximately 2 hours and was divided into twelve seven-minutes runs separated by eleven short breaks and one 30 min break.

### 2.3. MEG Data acquisition and preprocessing

MEG data were recorded using a whole-head 306 sensor (204 planar gradiometers; 102 magnetometers) Vector-view system (Elekta Neuromag, Helsinki, Finland). Five head-position indicator coils (HPIs) were used to continuously determine the head position with respect to the MEG helmet. MEG signals were recorded at a sampling rate of 1 kHz and an online band-pass filtered between 0.1 and 300 Hz. At the beginning of each experimental session, fiducial points of the head (the nasion and the left and right pre-auricular points) and a minimum of 300 other head-shape samples were digitized using a Polhemus FASTRAK 3D digitizer (Fastrak Polhemus, Inc., Colchester, VA, USA).

The raw data were processed using MaxFilter 2.0 (Elekta Neuromag ®). First, bad channels (identified via visual inspection) were replaced by interpolation. External sources of noise were separated from head-generated signals using a spatio-temporal variant of signal-space separation (SSS). Last, movement compensation was applied, and each run was aligned to an average head position. All further analysis steps were performed in MATLAB 2019a using non-commercial software packages such as Fieldtrip (Oostenveld et al., 2011), Brainstorm (Tadel et al., 2011) and custom scripts. Continuous MEG recordings were filtered at 0.1 Hz using a two-pass Butterworth high-pass filter and epoched from -1.5 s before to 2s after stimulus onset. Time segments contaminated by artifacts were manually rejected (total data loss of *M* = 2.4% *SD* = 1.8%). A Butterworth low-pass filter at 40Hz was applied to the epoched data. Before encoding, each trial segment was baseline corrected with respect to a -500 to -100ms time window before stimulus onset.

### 2.4. Multiple linear regression analysis

Multiple linear regression analysis was applied to MEG data following the approach used in previous M/EEG studies (Chen et al., 2013, 2015; Hauk et al., 2006, 2009; Miozzo et al., 2015). The solution of a multiple regression provides the best least-square fit of all variables simultaneously to the data (Bertero et al., 1985). For each time point, channel and subject we calculated event-related regression coefficients (ERRCs) reflecting the contribution of each predictor to the MEG signal. We focused on four predictors spanning word-form, lexical and semantic aspects of word retrieval (i.e., word length/duration, word frequency and an abstract and a concrete semantic predictor obtained via dimensionality reduction techniques of a 65 features’ space, see 2.5.). Before entering the regression model, regressors of interest (i.e., word length, word frequency, abstract semantic component and concrete semantic component) were orthogonalized via varimax rotation. Before encoding the predictors of each model were converted to normalized z-scores and tested for multicollinearity using a condition number test (Belsley, 1982). The output of the test is a condition index, which in the present study never exceeded a threshold of 2 (with test values < 6 collinearity is not seen as a problem).

### 2.5. Predictor variables

The aim of the present study was to investigate the contribution of abstract and concrete semantic dimensions of knowledge to concepts representations. On this account, we derived our stimulus set from a previous work by Binder and colleagues (Binder et al., 2016). These authors collected ratings of the salience of 65 biologically plausible features to word meaning (for a detailed description of the procedure see Binder et al. 2016). For every word in the database (e.g., lemon), more than one thousand participants were asked to rate how each of the features (e.g., color) was associated with that aspect of the experience (e.g., would you define a lemon as having a characteristic or defining color?). The result is a semantic space where concepts can be represented as single entities into a multidimensional space having perceptual (e.g., sound, shape, smell) and conceptual (e.g., arousal, social, sad) features as dimensions. Crucially, features spanned both abstract and concrete domains of conceptual knowledge thus represent an ideal framework to operationalize our assumptions.

#### 2.5.1. Semantic components

As mentioned above, more than sixty features composed our semantic space. Encoding the entire space in one single model, however, would be suboptimal. In fact, features are highly intercorrelated with each other, leaving us with a multicollinearity issue. One way this can be avoided is through dimensionality reduction techniques (Cunningham & Yu, 2014), such as principal component analysis (PCA). PCA generates a series of principal components (PCs) representing the same data in a new coordinate system, with the first PC usually accounting for the largest percentage of data variance. Following the concrete versus abstract labeling provided in the original database (Binder et al., 2016), we separated the entire semantic space (65-features) into concrete features (N = 31, encompassing Vision, Somatic, Audition Gustation, Olfaction and Motor domains) and abstract features (N = 31, encompassing Spatial, Temporal, Causal, Social, Emotion, Drive and Attention domains). Three features (i.e., Complexity, Practice, Caused) were excluded due to incomplete ratings. Thus, each word could be considered as a point in a concrete semantic features’ space (see Figure 1A), and in an abstract semantic features’ space (see Figure 1B). We used PCA to reduce the dimensionality of the dataset and adopted the first concrete semantic component (Figure 1C; 24.7% of variance explained) and the first abstract semantic component (Figure 1D; 27.4% of the variance explained), to represent the same data in a new one-dimensional coordinate system. Importantly, the resulting semantic components do not simply reflect how concrete and how abstract a word is, but instead represents concrete and abstract aspects of concepts in a new low-dimensional space that encodes the most salient structural features of the high-dimensional space from which it is derived. For instance, in the concrete principal component, “moose” is more similar to “street” than to “hug”, whereas the opposite is true in the abstract principal component (Figure 1, C-D).

**Figure 1.**
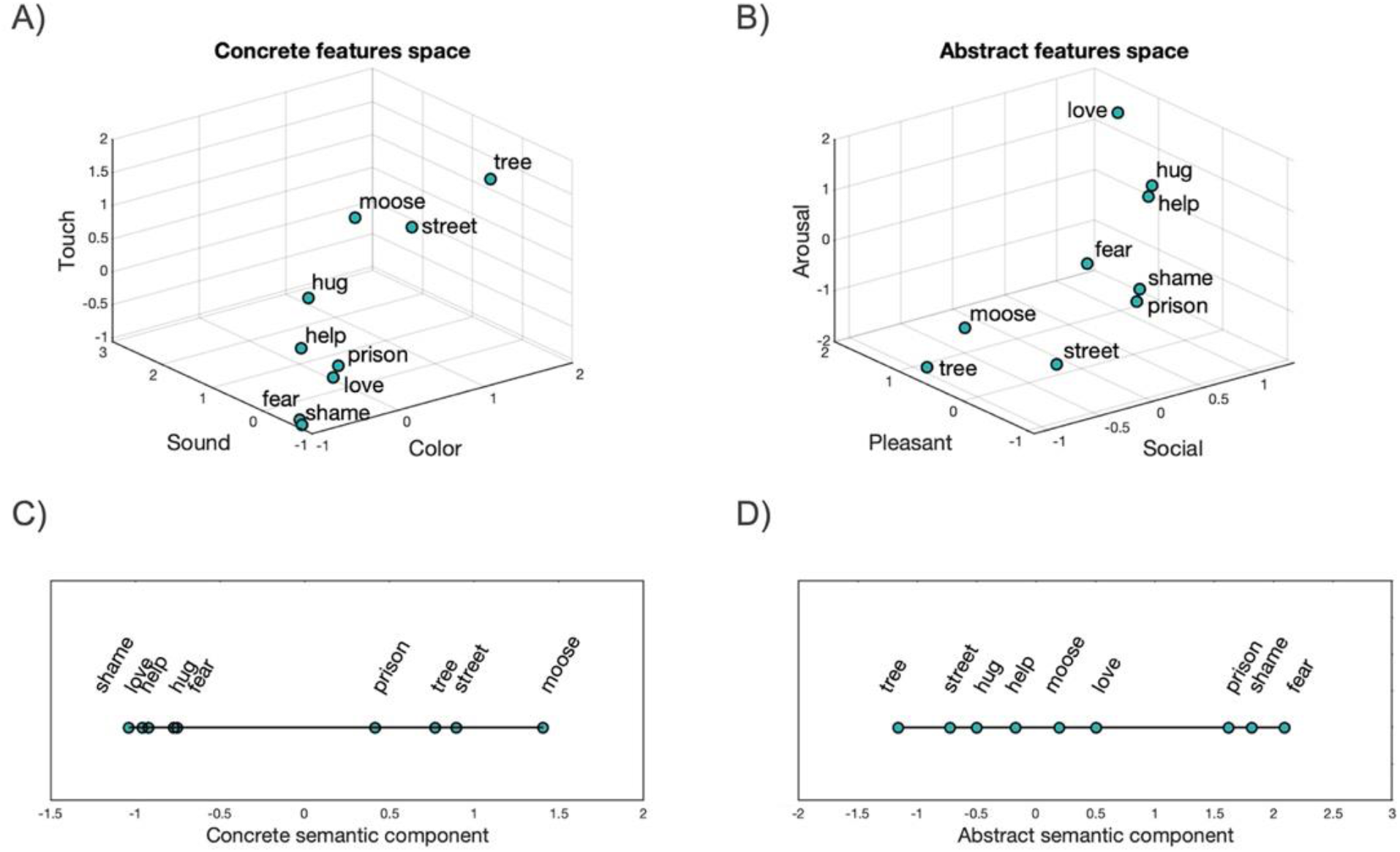
Dimensionality reduction. A) Schematic representation of a 3-D semantic space where each word is viewed in a coordinate system defined by concrete features such as Touch, Sound and Color (the actual multidimensional space comprised 31 dimensions, here reduced to 3 for visualization purposes). B) Schematic representation of a 3-D semantic space where each word is viewed in a coordinate system defined by abstract features such as Arousal, Pleasant and Social (the actual multidimensional space comprised 31 dimensions). C) Words’ weights along the first principal component of the concrete space. D) Words’ weights along the first principal component of the abstract space.

#### 2.5.2. Linguistic features

For each of the selected words, we obtained psycholinguistic features: Word Frequency (in Zipf’s scale, M = 4, SD = 0.8; van Heuven et al. 2014) was calculated as the frequency of occurrence of a given word in a large corpus of text samples (SUBTLEX-IT; Crepaldi et al. 2015). Word length was encoded as the number of letters of each word (*M* = 6.9, *SD* = 1.6).

### 2.6. Source reconstruction

Distributed minimum-norm source estimation (Hämäläinen & Ilmoniemi, 1994) was applied following the standard procedure in Brainstorm (Tadel et al., 2011). Anatomical T1-weighted MRI images were acquired during a separate session in a MAGNETOM Prisma 3T scanner (Siemens, Erlangen, Germany) using a 3D MPRAGE sequence, 1-mm^3^ resolution, TR = 2140ms, TI = 900ms, TE = 2.9ms, flip angle 12°. Anatomical MRI images were processed using an automated segmentation algorithm of the Freesurfer software (Fischl, 2012). Co-registration of MEG sensor configuration and the reconstructed scalp surfaces was based on ∼300 scalp surface locations. When no individual MRI was available (6 participants), we warped participants’ head shapes to a standard ICBM152 brain template. The data noise covariance matrix was calculated from the baseline interval (−500ms to -100ms) of the different trials. The forward model was obtained using the overlapping spheres method (Huang et al., 1999) as implemented in the Brainstorm software. We then: i) Estimated current density maps for event-related regression coefficients onto a 15000 vertices boundary element. Dipole sources were assumed to be perpendicular to the cortical surface. ii) Normalized current density values with respect to a -500ms to -100ms baseline period (z-transform). iii) Rectified current density values (converted to absolute values). iv) Spatially smoothed the source maps using an 8mm full width at half the maximum smoothing parameter (FWHM) and, finally, v) the individual results were projected to a default template (ICBM152).

### 2.7. Sensor-level statistical analysis and visualization

In line with previous studies (Chen et al., 2013, 2015; Hauk et al., 2006, 2009; Miozzo et al., 2015), we depicted the time course of different regressors as the root-mean-square (RMS) of the signal-to-noise ratio (SNR) of ERRC. The SNR was computed on the grand mean of all subjects by dividing the MEG signal at each channel and time point by the standard deviation of the baseline. This provided a unified (magnetometers and gradiometers are combined together) and easy-to-interpret measure of sensor-level activity. Statistical significance was assessed with t-test from -.5s to 1s after stimulus onset (FDR corrected for multiple comparisons, *p* < .05, Benjamini & Hochberg, 1995) on ERRC, separated for magnetometers and planar gradiometers (see Groppe et al., 2011). We additionally imposed temporal (a minimum duration of 20ms) as well as spatial (at least 2 concurrently significant channels) constraints on the reported results.

### 2.8. Source-level statistical analysis and visualization

Cortical responses to individual predictors (i.e., abstract semantic component, concrete semantic component, word frequency and word length; Figures 2 to 5, B) are illustrated as 20ms averages of source-reconstructed brain activity thresholded to the 80% of the local maxima. We additionally imposed temporal (a minimum duration of 20ms) as well as spatial (a minimum cluster size of 10 adjacent vertices) constraints on the reported results. Source-magnitude statistical maps (i.e., Concrete > Abstract, Figure 6) were computed using whole-brain t-tests (two-tailed), on consecutive 100ms average time windows (FDR corrected for multiple comparisons, *p* < .05, minimum number of 10 vertices).

**Figure 2.**
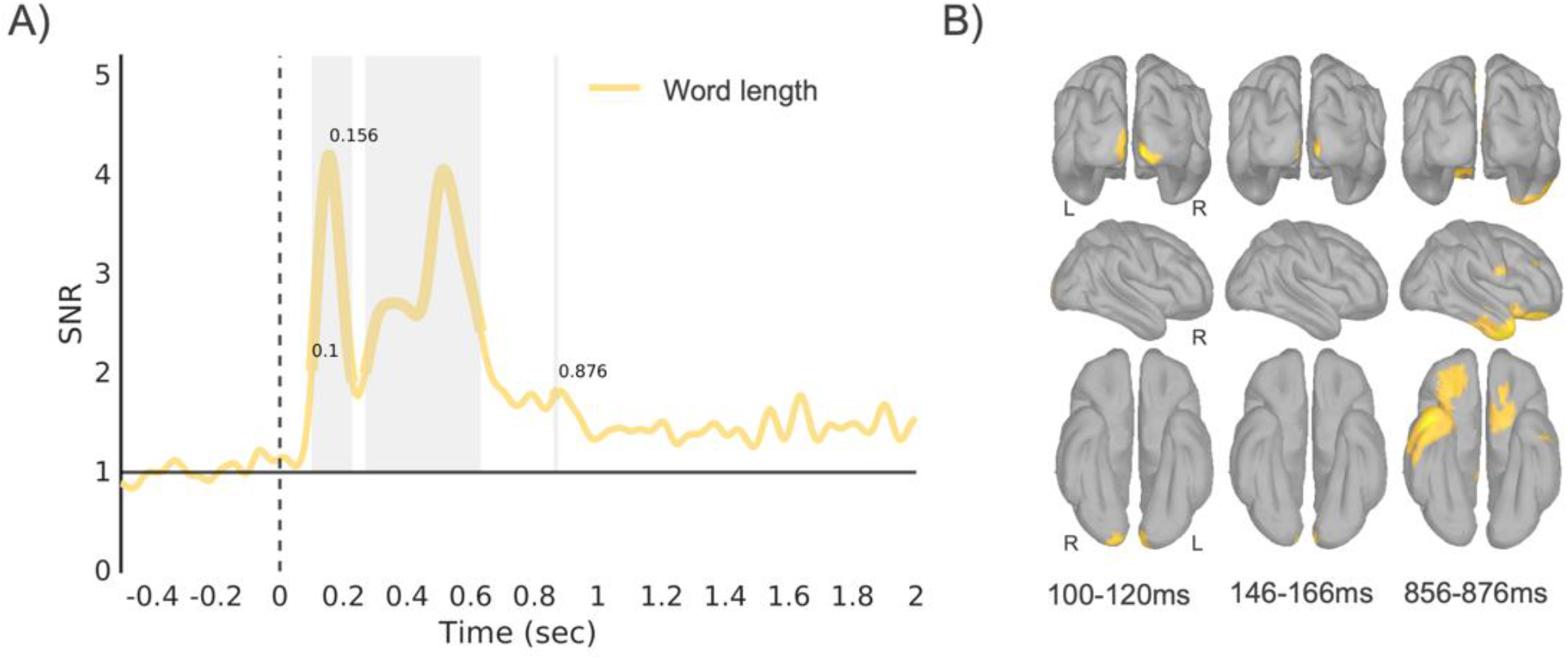
Spatiotemporal dynamics of word length information encoding. A) Sensor-level results depicted as the root-mean-square of the SNR of ERRC of the word length predictor. Significant time intervals (FDR corrected, p < .05) are indicated with a thicker line and a shadowed gray area. 0s = stimulus onset. B) Source-reconstructed maps of the word length predictor displayed as 20ms time averages (threshold 80% of local maxima, min cluster size 10, min duration 20ms) around the first significant time point (100-120ms), the peak of activation (146-166ms) and the last significant time point (856-876ms).

## 3. Results

### 3.1. Behavioral results

Participants were instructed to categorize each stimulus as either related to sensory perception (i.e., they refer to something that can be easily perceived with the senses, like “red” and “telephone”), or unrelated to sensory perception (i.e., they refer to something that cannot easily be perceived with the senses, like “agreement” and “shame”). We expected participants to categorize relatively concrete words as related to sensory perception and relatively abstract words as unrelated to sensory perception. To assess this, we correlated participants’ responses with the semantic principal components (see below). The results indicated a significant association between participants’ responses and our semantic dimensions (concrete semantic dimension: r(436)= .80, *p* < .001; abstract semantic dimension: r(436)= -.23, *p* < .001). We did not analyze reaction times because participants’ responses were delayed in order to avoid motion-related artifacts in the MEG signal (i.e., see Material and Methods for details).

### 3.2. Neural dynamics of lexical and semantic features

We first localized, in space and time, the encoding of the word length regressor (i.e., number of letters in a word). As predicted, this low-level visual information was encoded in and around primary visual cortices (bilaterally, Figure 2B), starting approximately 100ms after word appearance, peaking shortly after and remaining sustained up until 600ms after word onset (see Figure 2A). Such a highly predictable result served as a manipulation check for our source-localization procedure. At late time stages, word length information encoding saw the contribution of left inferior frontal and right anterior temporal and middle frontal brain systems.

Lexical access occurred shortly after processing of word-form related information. This is illustrated in Figure 3A, where encoding of the word frequency predictor (Zipf; van Heuven et al. 2014) begins around 300ms after visual word presentation, peaks at 580ms and continues until one second after stimulus onset. Source-level results are illustrated in Figure 3B. Encoding of information related to how frequent a word is in the language involves generators in the left ventral occipitotemporal cortex (approximately in the location of the Visual Word Form Area; Cohen et al. 2002) and anterior frontal brain regions. At the peak (approximately 600ms after word onset), these encompassed inferior frontal, anterior temporal, middle temporal and superior temporal brain areas with an overall moderate left lateralization. At later time points, the word frequency predictor was encoded in inferior frontal and anterior temporal brain areas, bilaterally (see Figure 3B).

**Figure 3.**
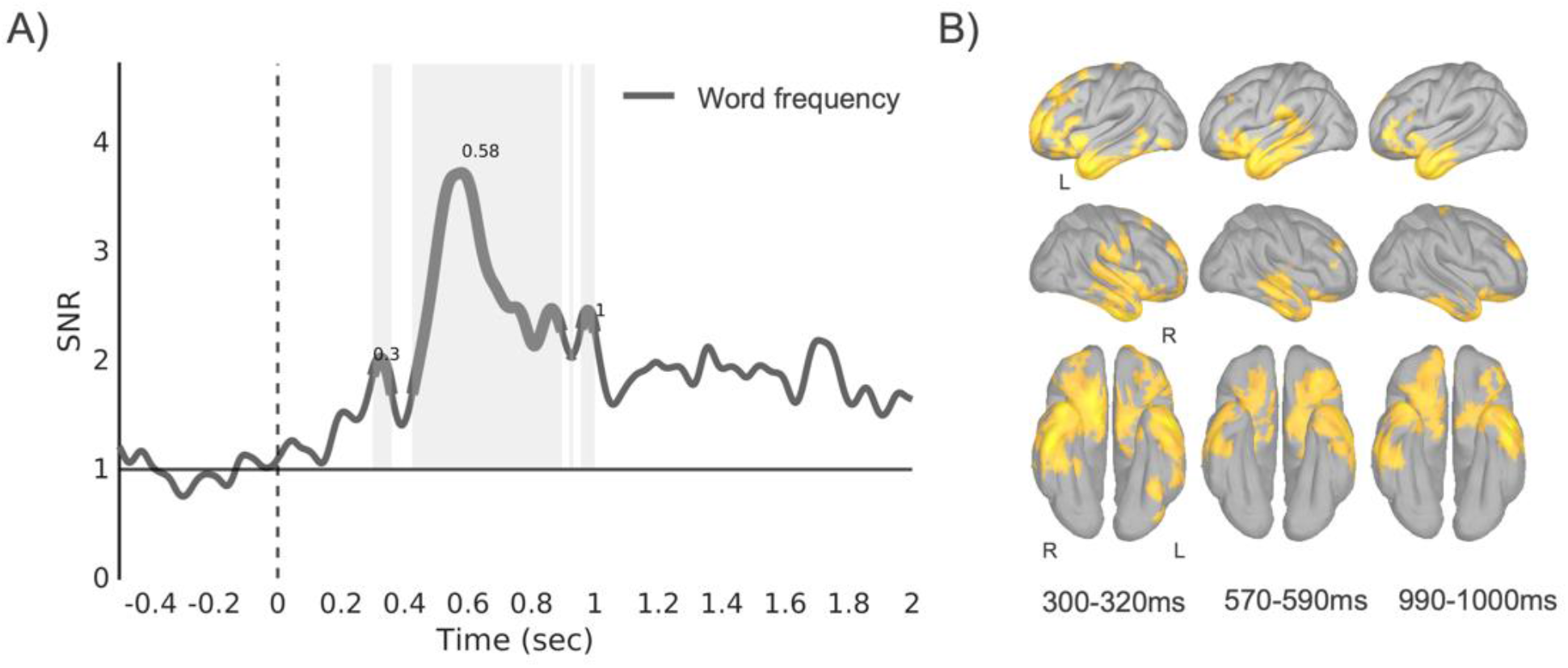
Spatiotemporal dynamics of word frequency information encoding. A) Sensor-level results depicted as the root-mean-square of the SNR of ERRC of the word frequency predictor. Significant time intervals (FDR corrected, p < .05) are indicated with a thicker line and a shadowed gray area. 0s = stimulus onset. B) Source-reconstructed maps of the word frequency predictor displayed as 20ms time averages (threshold 80% of local maxima, min cluster size 10, min duration 20ms) around the first significant time point (300-320ms), the peak of activation (570-590ms) and the last significant time point (990-1000ms).

Abstract semantic information processing began approximately 300ms after stimulus onset to peak 200ms after (see Figure 4A) and involved generators in prefrontal, inferior-frontal and anterior-temporal brain areas, bilaterally (see Figure 4B).

**Figure 4.**
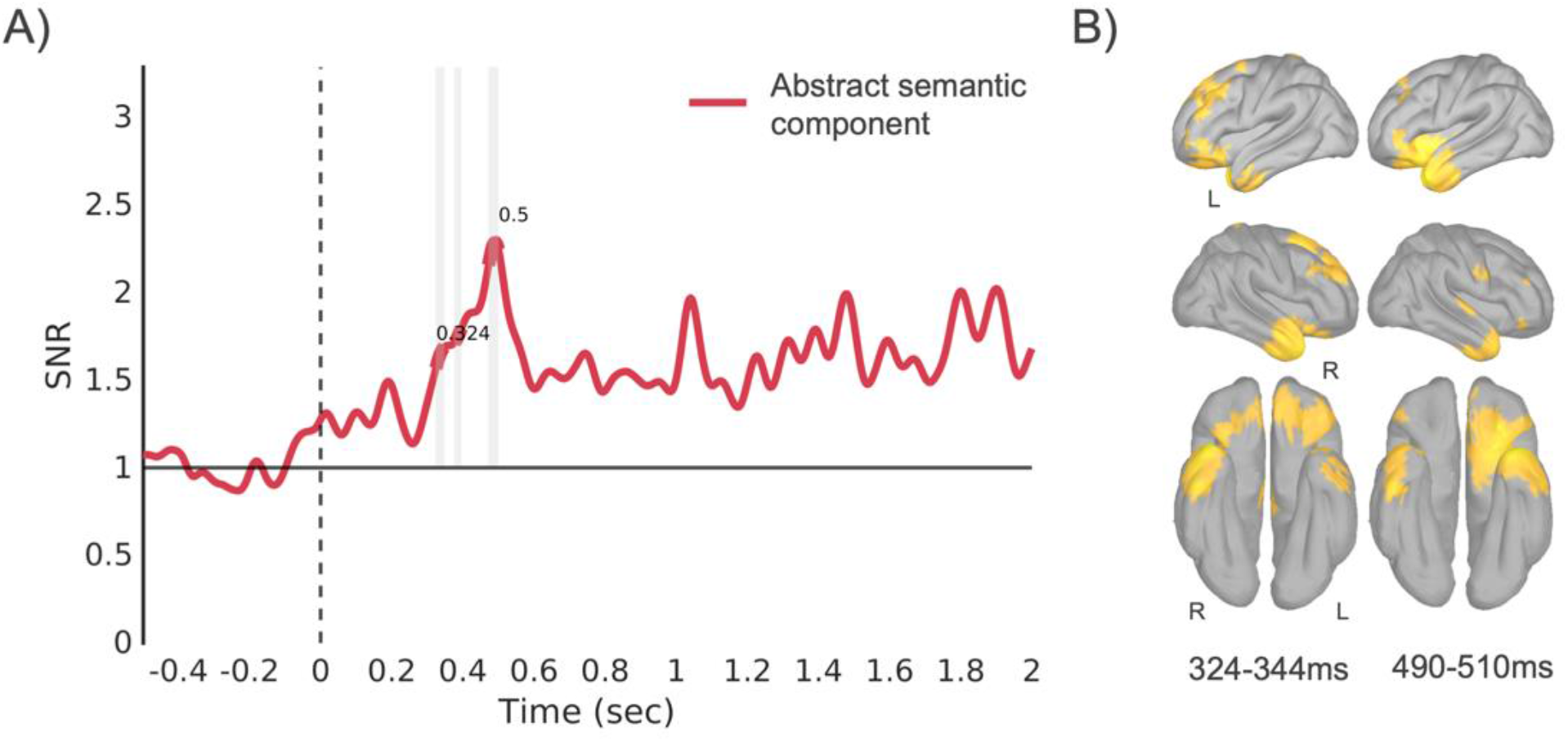
Spatiotemporal dynamics of abstract semantic information encoding. A) Sensor-level results depicted as the root-mean-square of the SNR of ERRC of the abstract semantic component. Significant time intervals (FDR corrected, p < .05) are indicated with a thicker line and a shadowed gray area. 0s = stimulus onset. B) Source-reconstructed maps of abstract semantic information encoding predictor displayed as 20ms time averages (threshold 80% of local maxima, min cluster size 10, min duration 20ms) around the first significant time point (324-344ms) and the peak of activation (490-510ms).

Encoding of concrete semantic information showed transient responses in the 300 to 500ms time window and a more sustained response from 600ms to 1s after stimulus onset (see Figure 5A). Source-level activation maps showed that concrete semantic information is encoded in prefrontal, inferior frontal and anterior temporal brain areas bilaterally (see Figure 5B).

**Figure 5.**
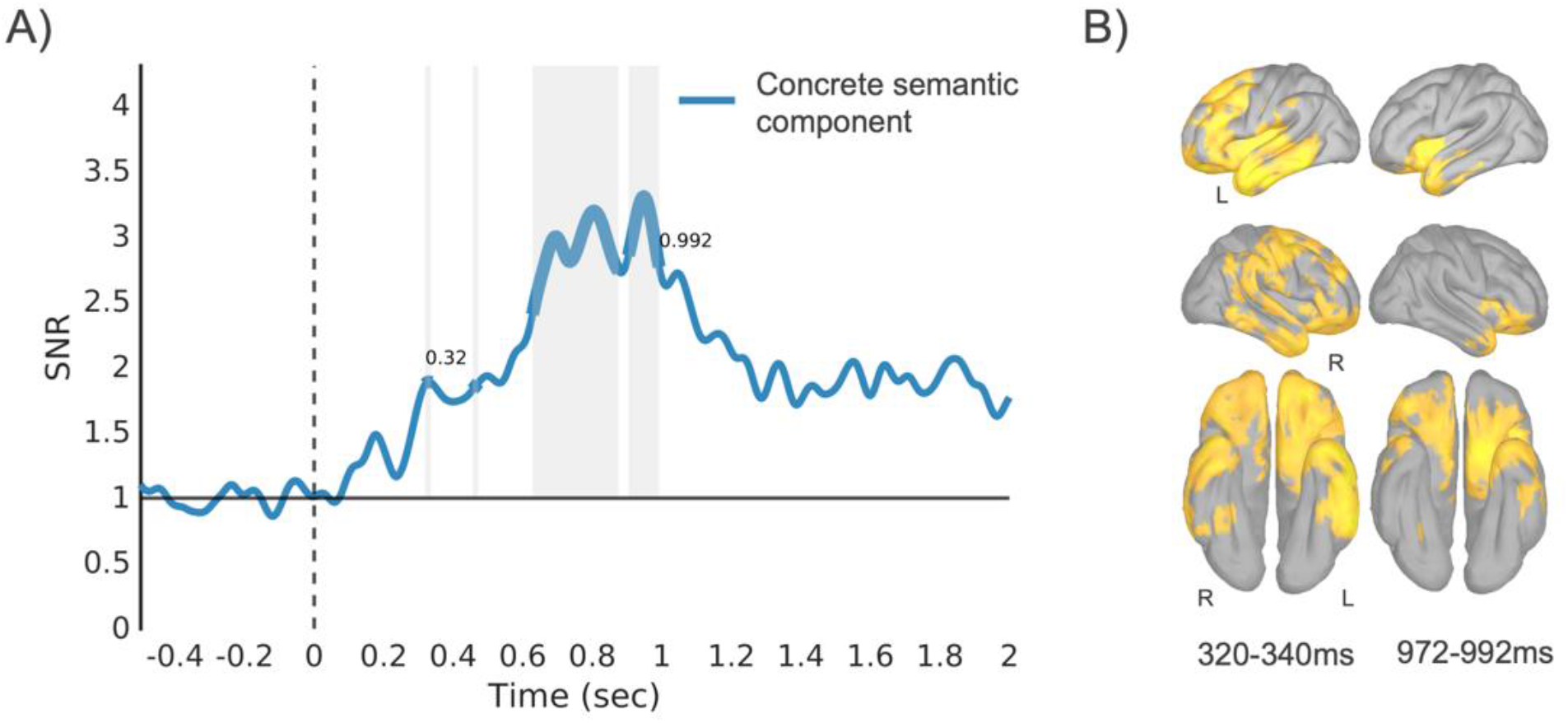
Spatiotemporal dynamics of concrete semantic information encoding. A) Sensor-level results depicted as the root-mean-square of the SNR of ERRC of the concrete semantic component. Significant time intervals (FDR corrected, p < .05) are indicated with a thicker line and a shadowed gray area. 0s = stimulus onset. B) Source-reconstructed maps of concrete semantic information encoding predictor displayed as 20ms time averages (threshold 80% of local maxima, min cluster size 10, min duration 20ms) around the first significant time point (320-340ms) and the last significant time point (972-992ms).

Last, we investigated source-magnitude activity of abstract and concrete semantic regressors which allowed us to describe in statistical terms brain areas showing greater responses to one or the other type of information. Results are illustrated in Figure 6 and evidenced greater activations for concrete semantic information in a distributed network of regions encompassing occipital, ventral occipito-temporal, inferior fusiform cortex and inferior-frontal brain areas approximately 700ms after word presentation. In line with a dual-coding approach, no brain region showed a greater activation for abstract compared to concrete features.

**Figure 6.**
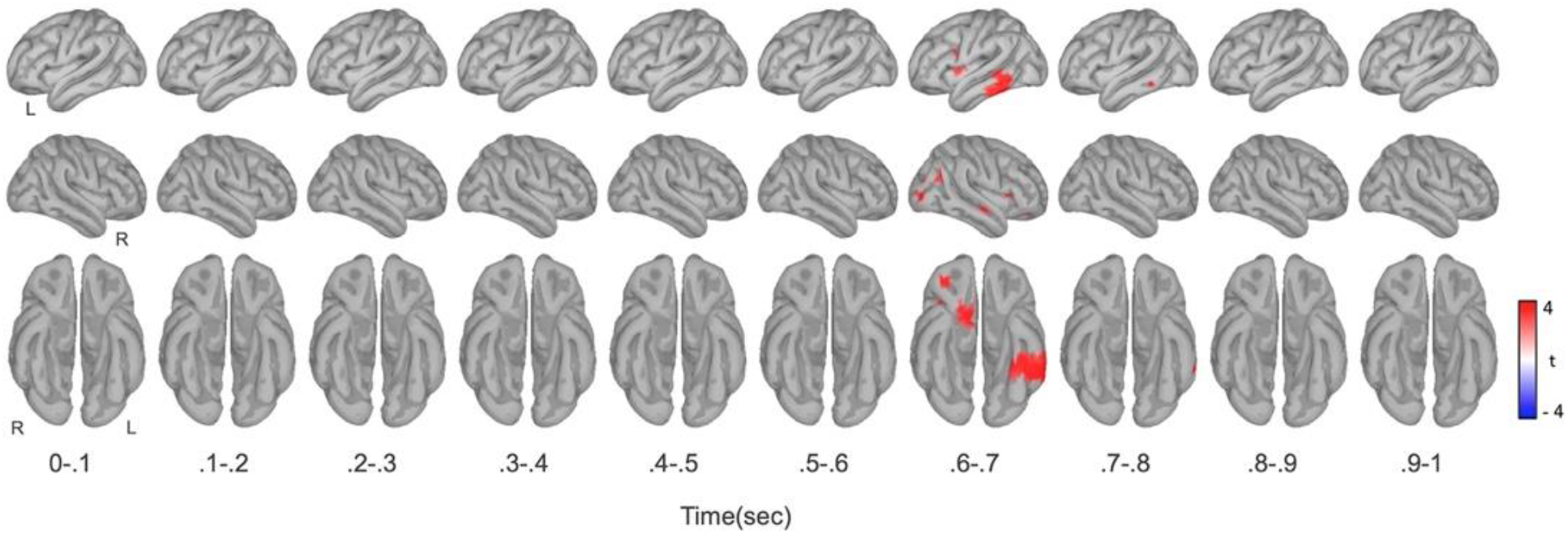
Analysis of source-level magnitude activations. Source-reconstructed statistical maps of the contrast Concrete > Abstract (paired-sample t-test (two-tailed), FDR-corrected *p* < .05, min cluster size 10) in consecutive 100ms average intervals.

## 4. Discussion

We took advantage of the high spatiotemporal resolution of MEG signals to test when and where abstract and concrete information is processed in the human brain. Using a multiple linear regression analysis of MEG-recorded brain activity, we obtained for every time point, channel, and subject event-related regression coefficients (ERRC) reflecting the contribution of each predictor to the data. Predictors of interest included variables associated with low-level visual information (the number of letters), lexical information (the frequency), as well as semantic properties (abstract and concrete feature dimensions) of each word.

Typically, the early stages of visual word recognition are dominated by the processing of low-level features (Carreiras et al., 2014). This is illustrated in Figure 2, where encoding of information related to the length of a word begins 100ms after stimulus onset and peaks shortly after (for similar findings, see Dufau et al., 2015; Hauk et al., 2009). Source analysis localized this result in bilateral occipital areas, reflecting the visual nature of these early contributions (see also Dhond et al., 2007; Hauk et al., 2009; Marinkovic et al., 2003). Sub-lexical information encoding was followed (∼200ms after) by lexico-semantic information encoding. That is, the word frequency predictor, the abstract semantic predictor, and the concrete semantic predictor all showed significant effects at around 300ms after stimulus onset (see Figures 3-4-5, A). The present findings reflect the cascade of underlying processes involved in visual word recognition (Grainger & Holcomb, 2009). A feedforward sweep of information cascades across sub-lexical and lexico-semantic stages resulting in parallel activations of lexical and semantic information approximately 300ms after word presentation (for similar findings, see Chen et al., 2015; Halgren et al., 2002; Pylkkänen & Marantz, 2003). At these latencies, the processing of information related to word frequency involved frontotemporal and left ventral occipitotemporal brain areas, consistent with functional imaging results of reading material (see, for instance, Kronbichler et al., 2004; Schurz et al., 2010; Schuster et al., 2016). Abstract and concrete semantic information processing, instead, involved a distributed network of brain areas encompassing both anterior frontal, anterior temporal and posterior brain areas (for similar findings, see Binder et al., 2009; Kana et al., 2012; Lambon-Ralph et al., 2017; Sabsevitz et al., 2005; Striem-Amit et al., 2018; J. Wang et al., 2010).

Recent dual coding accounts of knowledge suggest that meaning is represented in the brain by a double code, which comprises language-derived representations in the (dorsal) ATL and sensory-derived representations in perceptual and motor regions (Bi, 2021). This approach further predicts that anterior temporal regions should encode both concrete and abstract features, whereas perceptual and motor regions (e.g., occipital cortex) should encode mostly concrete features. Our results are in keeping with this view and provide additional information with respect to the temporal dynamics possibly underlying this cognitive model: As illustrated in Figures 4 and 5, encoding of both abstract and concrete semantic information showed early (300ms after word onset) engagement of anterior temporal and inferior frontal brain areas, suggesting that representations of word’s concreteness is not initially contingent on the activation of sensorimotor simulations or imagistic representations in perceptual and motor regions of the brain (Hauk et al., 2006; Hultén et al., 2021). Moreover, source-level analysis contrasting the abstract and concrete semantic regressors revealed that: (i) there was no brain region that was activated more by abstract compared to concrete features, as predicted by DCT; (ii) fusiform, lateral occipitotemporal, precentral and orbitofrontal regions preferentially encode concrete semantic features over abstract ones, in keeping with the prediction of DCT; (iii) finally, this neural signature emerged relatively late, around 700ms from word onset, suggesting a late activation of the sensorimotor/imagistic code during semantic processing.

Contrary to previous studies, a direct comparison of abstract and concrete semantic information encoding did not evidence stronger activations to abstract semantic information (see Figure 6). Whereas this result is in line with DCT, which predicts no differences between abstract and concrete representation in linguistic coding, greater activations to abstract concepts as to concrete concepts were reported in the linguistic areas in the inferior frontal cortex and the anterior temporal lobe (see, for instance, Binder et al., 2005; Hoffman et al., 2015). Our results did not confirm these observations, and this discrepancy might be due to task-induced mechanisms. Whereas our semantic categorization task (“Was it a word related to the senses? YES - NO”) put large emphasis on perceptual and motor representations of word meaning, several previous studies adopted tasks that emphasize linguistic properties of the stimuli (e.g., a lexical decision task, synonym judgment task; see for instance Binder et al., 2005; Hoffman et al., 2015). Wilson-Mendenhall et al. (2013) showed that under task conditions which require deeper conceptual processing, the linguistic system did not show stronger engagement with abstract concepts as compared to concrete concepts, in line with our results. It is possible that language-related tasks show greater sensitivity to symbolic/linguistic representations of abstract words, inducing a greater activation for such items in the language/symbolic network (Wilson-Mendenhall et al., 2013). However, it is also possible that lexical decision or synonym judgment foster the processing of lexical-semantic features such as semantic diversity (Hoffman et al., 2013), age of acquisition (Brown & Watson, 1987) or contextual availability (Schwanenflugel et al., 1988; Schwanenflugel & Stowe, 1989) which are often unbalanced between abstract and concrete words (abstract words usually have higher semantic diversity, lower contextual availability and are learned later in life). In this case, the higher activation of the language/symbolic network by abstract words could be due to the sensitivity of the network to such properties more than the preferential encoding of abstract semantic features per se.

Overall, the present findings suggest that contributions from a transmodal/linguistic code, housed in the perisylvian brain network, precede those of the imagistic/sensorimotor code in perceptual and motor regions, at least in the case of semantic concreteness. We cannot, however, exclude the prospect that, under different circumstances, this sequence of events would unfold differently. The case of action verbs may be a paradigmatic one, with many studies showing an early (∼200ms) activation of primary motor regions (e.g., M1) in response to action verbs (Hauk et al., 2008; Pulvermüller, 2013; Pulvermüller et al., 2005; van Elk et al., 2010). For instance, Garcia and colleagues (García et al., 2019), have recently shown that a machine learning classifier can distinguish action verbs (e.g., grasping) from nonaction verbs (e.g., sleeping), in M1, as early as 150ms after word onset. Interestingly, the same classification was found in ATL, but only later, around 250-300ms, thus revealing a reversed time course (sensorimotor regions before ATL) than the one we report here. Although their analyses were limited to these two regions of interest (ROIs), without control regions, and a limited number of participants, this data suggest that action verbs can activate simulations in primary motor regions during the very early stages of word processing (actually, as early as the peak of activation we found in primary visual cortex for word length; Figure 2).

However, taking into account the relevant exception of action verbs, in this experiment we showed that even when the task emphasized perceptual and motor representations of word meaning, posterior temporal, lateral occipital and precentral regions associated with a sensorimotor/imagistic code are preferentially activated by concrete features only during later stages of word processing. This finding supports the hypothesis that the concreteness advantage observed behaviorally during the early stages of word recognition can hardly be attributed to the activation of a sensorimotor/imagistic code in the sensorimotor regions of the brain (Bottini et al., 2021).

Dual code theories of knowledge in the brain successfully account for a large number of behavioral as well as neuroimaging findings (Bi, 2021; Paivio, 1986). Several aspects of this model, however, are still to be uncovered. It is for example unclear what is the exact nature of conceptual representations in the (dorsal) ATL? Is it really a language-based code that is “necessarily ‘amodal,’ ‘symbolic’ and independent from sensory experience” (Bi, 2021, p. 8)? In this view, the type of meaning supported by the linguistic code would be similar to the one encoded by current computational models in the field of natural language processing (NLP) and based on the statistical relationships with other words in speech (for a review Günther et al., 2019). In other words, the meaning is supported in language contexts (Barsalou et al., 2008; Vigliocco et al., 2009; Xu et al., 2017). However, this state of affairs begs the actual question behind the grounding problem (Harnad, 1990): If the linguistic code is ultimately granted by links between word forms, how can they entail meaning in the sense of referring to something beyond other words? Under this assumption, if congenitally blind people could rely only on the amodal, ungrounded and sensory independent linguistic code to understand the meaning of “red” they would find themselves trapped in the Chinese Room (Searle, 1980).

To solve this problem, several influential theories focus on the link between the two codes. One such example is “hub and spokes mode (H&S Patterson et al., 2007). H&S assumes that modality-specific sources of information (i.e., “spokes”), distributed across neocortical regions, encode different information sources (e.g., visual information in the occipital cortex, haptic in the sensorimotor cortex, linguistic in the perisylvian regions) that are integrated in the ATL “hub” (Lambon-Ralph et al., 2017; Patterson et al., 2007). In this model, the ATL is considered the home of transmodal representations that are not strictly language derived (language regions are one of the spokes in the model and simply one source of such integrated representations) but abstracted enough to affect all domains of knowledge (A. R. Damasio, 1989; H. Damasio et al., 1996; Patterson & Erzinçlioǧlu, 2008; Rogers & Patterson, 2007). Studies on functional connectivity corroborate this interpretation. By applying a graph-theoretic approach to the semantic brain network, Xu et al. (2017) highlighted two segregated systems for different types of semantic representations: a multimodal experiential content system in the default mode network and language supported content system in the perisylvian brain network. In this framework, anterior temporal areas are not the house to either linguistic or sensorimotor representations but are where these two representations converge (Xu et al., 2017). It has been also proposed that the ATL hub is organized according to a gradient of abstractness: The dorsal ATL would be more active for abstract concepts, given its preferential connectivity with perisylvian language regions; Whereas the ventro-medial ATL would be more active for concrete concepts given its connections with visual brain regions. However, the spatial resolution of MEG is limited and makes the distinction between subparts of the ATL difficult to achieve. Finally, our design, does not allow to disentangle whether linguistic, or integrated representations (or both) encoded abstract and concrete semantic features in anterior temporal regions.

## 5. Conclusions

To conclude, the present findings shed new light on the spatiotemporal dynamics of abstract and concrete semantic representations in the brain. At early processing stages, abstract and concrete semantic information encoding was underpinned by common neural substrates in the anterior temporal lobe, whereas at later latencies, sensory-motor areas showed preferential responses to concrete information only. We suggest that concreteness is encoded in the brain via the early contribution of a transmodal/linguistic code (housed in frontotemporal brain systems), followed by the activation of an imagistic/sensorimotor code in perceptual regions. Results are broadly consistent with a dual-coding approach, although the strictly linguistic nature of ATL representations remains putative and waits for further empirical studies.

## Author contributions

**L. Vignali:** Conceptualization, Data curation, Formal analysis, Investigation, Software, Methodology, Visualization, Writing - Original draft. **Y. Xu:** Conceptualization, Formal analysis, Writing – Review & Editing. **J. Turini:** Investigation, Resources, Data curation. **O. Collignon:** Conceptualization, Writing – Review & Editing, Supervision, Funding acquisition. **D. Crepaldi:** Conceptualization, Writing – Review & Editing, Supervision, Funding acquisition. **R. Bottini:** Conceptualization, Writing – Review & Editing, Supervision, Project administration, Funding acquisition.

## Declaration of competing interest

The authors declare that they have no known competing financial interests or personal relationships that could have appeared to influence the work reported in this paper.

## Acknowledgments

The project was partly funded by a PRIN grant (Project number: 2015PCNJ5F_001) from the Italian Ministry of Education, University and Research (MIUR) awarded to Davide Crepaldi in collaboration with Olivier Collignon and by an European Research Council grant (H2020 ERC G.A. n: 804422) awarded to Roberto Bottini.

## Notes

### Competing Interest Statement

The authors have declared no competing interest.

